# Placental expression of ACE2 and TMPRSS2 in maternal SARS-CoV-2 infection: are placental defenses mediated by fetal sex?

**DOI:** 10.1101/2021.04.01.438089

**Authors:** Lydia L Shook, Evan A Bordt, Marie-Charlotte Meinsohn, David Pepin, Rose M De Guzman, Sara Brigida, Laura J Yockey, Kaitlyn E James, Mackenzie W Sullivan, Lisa M Bebell, Drucilla J Roberts, Anjali J Kaimal, Jonathan Z Li, Danny Schust, Kathryn J Gray, Andrea G Edlow

## Abstract

**Background:** Sex differences in vulnerability to and severity of SARS-CoV-2 infection have been described in non-pregnant populations. ACE2 and TMPRSS2, host molecules required for viral entry, are regulated by sex steroids and expressed in the placenta. We sought to investigate whether placental *ACE2* and *TMPRSS2* expression vary by fetal sex and in the presence of maternal SARS-CoV-2 infection.

**Methods:** Placental ACE2 and TMPRSS2 were quantified in 68 pregnant individuals (38 SARS-CoV-2 positive, 30 SARS-CoV-2 negative) delivering at Mass General Brigham from April to June 2020. Maternal SARS-CoV-2 status was determined by nasopharyngeal RT-PCR. Placental SARS-CoV-2 viral load was quantified. RTqPCR was performed to quantify expression of *ACE2* and *TMPRSS2* relative to the reference gene *YWHAZ*. Western blots were performed on placental homogenates to quantify protein levels. The impact of fetal sex and SARS-CoV-2 exposure on ACE2 and TMPRSS2 expression was analyzed by 2-way ANOVA.

**Results:** SARS-CoV-2 virus was undetectable in all placentas. Maternal SARS-CoV-2 infection impacted TMPRSS2 placental gene and protein expression in a sexually dimorphic fashion (2-way ANOVA interaction p-value: 0.002). We observed no impact of fetal sex or maternal SARS-CoV-2 status on placental ACE2 gene or protein expression. Placental *TMPRSS2* expression was significantly correlated with *ACE2* expression in males (Spearman’s ρ=0.54, p=0.02) but not females (ρ=0.23, p=0.34) exposed to maternal SARS-CoV-2.

**Conclusions:** Sex differences in placental TMPRSS2 but not ACE2 were observed in the setting of maternal SARS-CoV-2 infection. These findings may have implications for offspring vulnerability to placental infection and vertical transmission.These findings may have implications for offspring vulnerability to placental infection and vertical transmission.

## Background

More than 82,000 pregnant women in the United States have tested positive for SARS-CoV-2 [1], with numbers rising daily. Reported rates of vertical transmission of SARS-CoV-2 – the passage of SARS-CoV-2 from mother to baby during pregnancy or childbirth – range from 1-3% [2–4]. Case reports and case series documenting placental SARS-CoV-2 infection in the second and third trimesters suggest that SARS-CoV-2 infection of the placenta may be an intermediate step in vertical transmission [5–11]. The relative rarity of placental SARS-CoV-2 infection and vertical transmission may be driven by a low incidence of maternal viremia, and protective patterns of SARS-CoV-2 entry receptors within the placenta [12].

Infection of host cells by SARS-CoV-2, the novel coronavirus responsible for COVID-19, requires the presence of two molecules: the transmembrane receptor Angiotensin-Converting Enzyme 2 (ACE2) and the Type II Transmembrane Serine Protease (TMPRSS2) [13,14]. TMPRSS2 primes the SARS-CoV-2 Spike protein which subsequently binds to ACE2 as the point of entry into the host cell [14]. Immunohistochemical analyses in placentas exposed to maternal SARS-CoV-2 demonstrate strong expression of ACE2 in syncytiotrophoblast and cytotrophoblast across gestation, but weak expression of TMPRSS2, most often confined to the fetal endothelium or unquantifiable by conventional immunohistochemical methods [12,15,16]. Single cell placental atlases from samples predating the COVID-19 pandemic demonstrate single cell gene expression of *ACE2*, but offer conflicting results regarding co-transcription of *TMPRSS2*, and co-expression of *ACE2* and *TMPRSS2* within the same cell type at the maternal-fetal interface [17–20]. There is therefore a lack of information on how maternal infection with SARS-CoV-2 impacts the expression of *ACE2* and *TMPRSS2* in the placenta, at the level of both transcript and protein.

There is an urgent need to identify factors, including biological sex, that may influence ACE2 and TMPRSS2 expression and functionality in human tissues. Epidemiologic data point to a male bias in susceptibility to severe COVID-19 disease and mortality [21–25], with sex differences observed in outcomes across the lifespan [25–28]. More severe disease in male infants and children has been demonstrated in population level and cohort data reporting a male predominance in children with multisystem inflammatory syndrome in children (MIS-C), a severe form of COVID-19 disease in children, and in reported infections in the age range from birth to two years [27–31]. Sex differences in the expression of ACE2 and TMPRSS2 in respiratory and cardiovascular tissues has been hypothesized to underlie these population-level observations [25,32–36]. Examining sex differences in placental expression of ACE2 and TMPRSS2 in the setting of maternal SARS-CoV-2 infection has the potential to yield insights into placental defenses against vertical transmission.

Whether male offspring are more susceptible to vertical transmission is not well-characterized, as vertical transmission is relatively uncommon, and sex disaggregated data are not routinely reported [2,3]. Given available data supporting a connection between COVID-19 outcomes and sex differences in ACE2 and TMPRSS2 expression levels [32,36–38], and the paucity of information on ACE2 and TMPRSS2 expression during pregnancy, we sought to investigate whether placental expression of ACE2 and TMPRSS2 varies by fetal sex in the presence and absence of maternal SARS-CoV-2 infection.

## Methods

### Study design and participant enrollment

These experiments include 68 pregnant women (38 SARS-CoV-2 positive, 30 SARS-CoV-2 negative) enrolled in a cohort study at Massachusetts General Hospital and Brigham and Women’s Hospital from April to June 2020, coincident with universal screening for SARS-CoV-2 infection by nasopharyngeal RT-PCR on the Labor and Delivery unit. This study was approved by the Mass General Brigham Institutional Review Board (#2020P003538). All participants provided written informed consent.

Pregnant women were eligible for inclusion if they were: (1) 18 years of age or older, (2) able to provide informed consent or had a named healthcare proxy to do so, and (3) diagnosed with SARS-CoV-2 infection or known to be negative for SARS-CoV-2 by nasopharyngeal swab RT-PCR. Maternal SARS-CoV-2 positivity was defined as a positive nasopharyngeal swab RT-PCR at any time during pregnancy. Identification of eligible women and participant enrollment have been described in previous publications [12,39] and are summarized here. Participants positive for SARS-CoV-2 were identified either on admission to the Labor and Delivery unit through universal SARS-CoV-2 screening (initiated early April 2020) by nasopharyngeal RT-PCR, or by a prior documented positive SARS-CoV-2 nasopharyngeal RT-PCR during pregnancy for a COVID-19-related illness. Participants negative for SARS-CoV-2 on admission to Labor and Delivery were enrolled as a convenience sample, recruited on the same days as enrolled positive cases. Demographic and clinical outcomes data were abstracted from the electronic medical record using REDCap electronic data capture tools [40]. Included participants were selected to balance groups for fetal sex, and to distribute maternal disease severity evenly across fetal sex categories. COVID-19 disease severity was defined according to NIH criteria [41].

### Sample collection and processing

Sample collection protocols have been described in a previous publication [39]. Briefly, at the time of delivery, two ∼0.5 cm^3^ placental biopsies were collected from the maternal side of the placenta and two from the fetal side of the placenta, at least 4 cm from the cord insertion and the placental edge, after dissecting off the overlying amnion and chorion or decidua. Biopsies were placed in 5 mL RNALater and stored at 4°C for at least 24 hours. Biopsies were then flash frozen and stored at -80°C.

### RNA extraction

To control for regional differences in placental gene expression, two maternal side and two fetal side placental samples (∼50 mg tissue per side) were processed per placenta. Total RNA was obtained following homogenization of tissue sample in Trizol (100 uL/10 mg). Resulting suspension was then centrifuged at 12,000 x g for 10 min after which the pellet was discarded. Chloroform was added to supernatant at a ratio of 100 uL chloroform/50 mg tissue. Tubes were shaken vigorously for 15 seconds, allowed to stand at room temperature for 10 min, and then centrifuged at 12,000 x g for 15 min. The aqueous phase was collected and the remainder of the RNA extraction procedure was performed using an RNeasy Mini Kit with on-column DNase I treatment (Qiagen) according to manufacturer instructions. RNA quantity and purity were assessed using a NanoDrop 2000 Spectrophotometer (ThermoFisher Scientific).

### cDNA synthesis and RTqPCR

cDNA synthesis was performed using iScriptTM cDNA Synthesis Kit (Bio-Rad) per manufacturer instructions using a MiniAmpTM Plus Thermal Cycler (ThermoFisher Scientific). No template control and no RT controls were prepared. Quantitative real-time polymerase chain reaction (RTqPCR) was then performed using Taqman gene expression assays on a QuantStudio 5 Real-Time PCR System (ThermoFisher Scientific). Gene expression was normalized to the placental reference gene *Tyrosine 3-Monooxygenase/Tryptophan 5-Monooxygenase Activation Protein Zeta* (*YWHAZ*) and expressed relative to female SARS-CoV-2 negative control samples to yield a relative quantity value (2^-ΔΔCt^). *YWHAZ* was selected due to its prior validation as a stably-expressed reference gene in placental tissue [42–44], and no changes in *YWHAZ* expression were noted by fetal sex or maternal SARS-CoV-2 infection Gene expression assays used include *ACE2* Hs01085333_m1, *TMPRSS2* Hs01122322_m1, and *YWHAZ* Hs01122445_g1 (TaqMan, ThermoFisher Scientific). *ACE2* and *TMPRSS2* TaqMan gene expression assays used FAM-MGB dye, the *YWHAZ* assay used VIC_PL dye. Cases and controls were run on the same plates using the same mastermix for all samples, and reactions were performed in triplicate.

### Quantification of SARS-CoV-2 viral load by RT-PCR

SARS-CoV-2 viral load was quantified from extracted placental RNA using the US CDC 2019-nCoV_N1 primers and probe set [45]. Viral copy numbers were quantified using N1 qPCR standards in 16-fold dilutions to generate a standard curve. The assay was run in triplicate for each sample. Positive controls, negative controls, and two non-template control (NTC) wells were included as negative controls. Quantification of *glyceraldehyde-3-phosphate dehydrogenase* (*GAPDH*) reference gene RNA level was performed to determine the efficiency of RNA extraction and qPCR amplification (Bio-Rad PrimePCR assay #qHsaCED0038674). SARS-CoV-2 viral loads below 40 RNA copies/mL were categorized as undetectable and set at 1.0 log_10_ RNA copies/mL.

### Western blots

56 cases (28 female, 28 male) were examined for expression of ACE2 and TMPRSS2 by Western blot. 30 ug of protein was prepared in SDS-PAGE Sample Loading Buffer (G Biosciences) with 5 mM dithiothreitol (DTT) and heated for 10 min at 80°C, and loaded on Mini-PROTEAN 10% TGX Stain-Free Protein Gels (Bio-Rad). Gels were run at 200V for 30 min, after which they were transferred onto low fluorescence PVDF membrane using a Trans-Blot Turbo Transfer System (Bio-Rad). Blots were then washed 3x 10 min in TBS followed by imaging of Stain Free total protein signal with a ChemiDoc MP System (Bio-Rad). Following stain-free imaging, blots were washed another 3 x 10 min in TBS, followed by blocking in Intercept (TBS) Blocking Buffer (Li-Cor Biosciences) for 1 hr at room temperature. ACE2 and TMPRSS2 primary antibodies were then incubated in Intercept T20 (TBS) Antibody Diluent (Li-Cor Biosciences) overnight at 4°C. Membranes were then washed 6 x 10 min in TBS-T, followed by incubation with secondary antibodies at 1:2500 dilution for 1 hr in Intercept T20 (TBS) Antibody Diluent. Primary and secondary antibodies, manufacturer, and dilution conditions used for immunoblot are detailed in Table 1. Blots were washed 6 x 10 min in TBS-T, after which they were briefly rinsed in TBS prior to imaging. Band intensity was then imaged using a ChemiDoc MP System and quantified using Image Lab Software (Bio-Rad). Data are presented as the volume of bands of protein of interest relative to the volume of total protein quantified using stain-free imaging.

**Table 1.** Primary and secondary antibodies, manufacturer, and dilution conditions used for Western blot.

| **Antibody** | **Manufacturer** | **Catalog #** | **Dilution** |
| --- | --- | --- | --- |
| ACE2 antibody | Abcam | ab108252 | 1:500 |
| TMPRSS2 | Novus | NBP1-20984 | 1:2,500 |
| Donkey anti-rabbit (HRP) | Abcam | ab205722 | 1:2,500 |
| Donkey anti-goat (HRP) | ThermoFisher | A15999 | 1:2,500 |

### Statistical analysis

All data are presented as mean ± standard error of the mean in the text and Figure Legends unless otherwise noted. Relative gene expression values (relative to *YWHAZ*) are expressed as arbitrary units. Two-way analysis of variance (2-way ANOVA) with Bonferroni’s post-hoc testing was used to compare gene expression by fetal sex and maternal SARS-CoV-2 infection, and Western immunoblot staining intensity by fetal sex and maternal SARS-CoV-2 infection. Correlations between *ACE2* and *TMPRSS2* mRNA levels were assessed using Spearman’s rank-order correlation. P < 0.05 was considered statistically significant. Statistical analyses were performed using GraphPad Prism 9.

## Results

### Participant characteristics

Maternal demographic and clinical data of study participants are depicted in Table 2. Of the 68 included participants, 34 were pregnant with females and 34 with males, with 19 SARS-CoV-2 positive and 15 SARS-CoV-2 negative in each group. There were no significant differences in maternal age, gravidity, parity, race, or ethnicity. Of the 38 women with SARS-CoV-2 infection during pregnancy, there were no differences between male and female pregnancies with respect to gestational age at diagnosis of SARS-CoV-2 infection, days from SARS-CoV-2 diagnosis to delivery, or severity of COVID-19 illness. Women with SARS-CoV-2 infection during pregnancy were more likely to be Hispanic compared to uninfected controls, consistent with prior report of ethnic disparities in COVID-19 vulnerability in our patient population [46].

**Table 2.** Demographic and clinical characteristics of pregnancy cohort by fetal sex and maternal SARS-CoV-2 status. Abbreviations: BMI, body mass index; GA, gestational age; N/A, not applicable. Continuous variables presented as median [IQR]. ^a^”SARS-CoV-2 positive” indicates positive test for SARS-CoV-2 by nasopharyngeal RT-PCR during pregnancy ^b^Significant differences between groups were determined using chi-square test for categorical variables, and Kruskall-Wallis test for continuous variables. ^c^”Hypertension” indicates chronic or pregnancy-associated hypertension ^d^”Diabetes” indicates pre-existing or gestational diabetes ^e^Severity determinations were made at the time of diagnosis and based on published criteria from the National Institutes of Health

|  | All (68) | Female |  | Male |  | P<sup>b</sup> |
| --- | --- | --- | --- | --- | --- | --- |
|  |  | SARS-CoV-2<br>negative<br>(15) | SARS-CoV-2<br>positive<br>(19)<sup>a</sup> | SARS-CoV-2<br>negative<br>(15) | SARS-CoV-2<br>positive<br>(19)<sup>a</sup> |  |
| Maternal<br>Age, years | 33<br>[28-38] | 33<br>[30-36] | 34<br>[30-40] | 30<br>[27-37] | 31<br>[26-36] | 0.34 |
| Parity, median<br>[IQR] | 1<br>[0-2] | 1<br>[0-2] | 1<br>[0-2] | 1<br>[0-2] | 1<br>[0-2] | 0.34 |
| Race, n (%) |  |  |  |  |  | 0.19 |
| White | 40 (59) | 11 (73) | 8 (42) | 12 (80) | 9 (59) |  |
| Black | 5 (7) | 1 (7) | 3 (16) | 0 (0) | 1 (7) |  |
| Asian | 1 (1) | 0 (0) | 0 (0) | 1 (7) | 0 (0) |  |
| Other | 12 (18) | 3 (20) | 4 (21) | 1 (7) | 4 (8) |  |
| Not Reported | 10 (15) | 0 (0) | 4 (21) | 1 (7) | 5 (15) |  |
| Ethnicity, n (%) |  |  |  |  |  | 0.01 |
| Hispanic | 32 (47) | 4 (27) | 11 (58) | 2 (13) | 15 (79) |  |
| Non-Hispanic | 33 (48) | 10 (67) | 7 (37) | 12 (80) | 4 (21) |  |
| Not Reported | 3 (4) | 1 (7) | 1 (5) | 1 (7) | 0 (0) |  |
| Hypertension<sup>c</sup>, n<br>(%) | 8 (12) | 1(7) | 1 (5) | 2 (13) | 4 (21) | 0.43 |
| Diabetes<sup>d</sup>, n (%) | 6 (9) | 1 (7) | 3 (16) | 1 (7) | 1 (5) | 0.67 |
| BMI ≥ 30 kg/m<sup>2</sup>, n<br>(%) | 20 (29) | 2 (13) | 7 (37) | 3 (20) | 8 (42) | 0.21 |
| Pre-pregnancy<br>BMI, kg/m<sup>2</sup><br>(median [IQR]) | 27.4<br>[21.5-30.8] | 25.8<br>[21.6-29.4] | 28.5<br>[26.5-32.7] | 21.5<br>[20.3-29.8] | 29.5<br>[22.3-32.0] | 0.06 |
| GA at<br>delivery, weeks<br>(median [IQR]) | 39.2<br>[38.6-40.3] | 39.3<br>[39.0-39.4] | 39.3<br>[38.4-40.1] | 39.1<br>[39.0-41.1] | 39.0<br>[35.3-40.3] | 0.09 |
| Any labor, n (%) | 52 (76) | 8 (53) | 16 (84) | 11 (73) | 17 (89) | 0.07 |
| Neonatal birthweight, g (median [IQR]) | 3283<br>[3005-3590] | 3400<br>[3060-3590] | 3255<br>[2920-3340] | 3435<br>[3260-3730] | 3115<br>[2615-3590] | 0.06 |
| GA at positive SARS-CoV-2 test, weeks (median [IQR]) | 38.7<br>[35.6-39.6] | N/A | 36.3<br>[32.4-39.4] | N/A | 36.3<br>[32.4-39.5] | 0.90 |
| COVID-19 disease severity<sup>e</sup>, n (%) |  |  |  |  |  | 0.82 |
| *Asymptomatic* | 16 (42) | N/A | 8 (42) | N/A | 8 (42) |  |
| *Mild/Moderate* | 19 (50) | N/A | 9 (47) | N/A | 10 (53) |  |
| *Severe/Critical* | 3 (8) | N/A | 2 (11) | N/A | 1 (5) |  |
| Time between SARS-CoV-2 symptom onset and delivery, days (median [IQR]) | 36.5<br>[18-57] | N/A | 44 [28-64] | N/A | 32 [8-51] | 0.18 |

### Placental SARS-CoV-2 viral load

SARS-CoV-2 viral load was undetectable (1.0 log_10_ RNA copies/mL) in all 68 placentas, consistent with our previous findings demonstrating no cases of SARS-CoV-2 RNA in the placenta by RNA *in situ* hybridization [12]. The same viral load assay was used to detect SARS-CoV-2 in respiratory samples from pregnant study participants and blood from non-pregnant study participants [12,47].

### TMPRSS2 placental expression was impacted by fetal sex and maternal SARS-CoV-2 exposure

As co-expression of ACE2 and TMPRSS2 is critical for viral entry to host cells [13], we sought to determine whether *TMPRSS2* levels were altered by SARS-CoV-2 exposure or by fetal sex. We found that maternal exposure to SARS-CoV-2 impacted *TMPRSS2* placental gene expression in a sexually dimorphic fashion (2-way ANOVA, fetal sex: P=0.06, maternal SARS-CoV-2 P=0.35, interaction: P=0.002, Figure 1A). *TMPRSS2* expression was significantly higher in male compared to female placentas of participants negative for SARS-CoV-2 (adjusted P=0.006). *TMPRSS2* expression was significantly reduced in male placentas of participants positive for SARS-CoV-2 compared to male placentas of SARS-CoV-2 negative controls (adjusted P=0.02, Figure 1A). The same pattern was observed at the protein level by Western blot (2-way ANOVA, fetal sex: P=0.65, maternal SARS-CoV-2 P=0.90, interaction: P=0.0002, Figure 1B-C). TMPRSS2 protein levels were significantly elevated in male controls compared to female controls (adjusted P=0.01). A sexually dimorphic response to maternal SARS-CoV-2 infection was observed, in which TMPRSS2 levels were increased in females (adjusted P=0.03) but decreased in males (adjusted P=0.04) exposed to maternal SARS-CoV-2 infection, relative to sex-matched controls. Table 3 depicts the results of 2-way ANOVA analyses of qPCR and Western blot results.

**Table 3.**
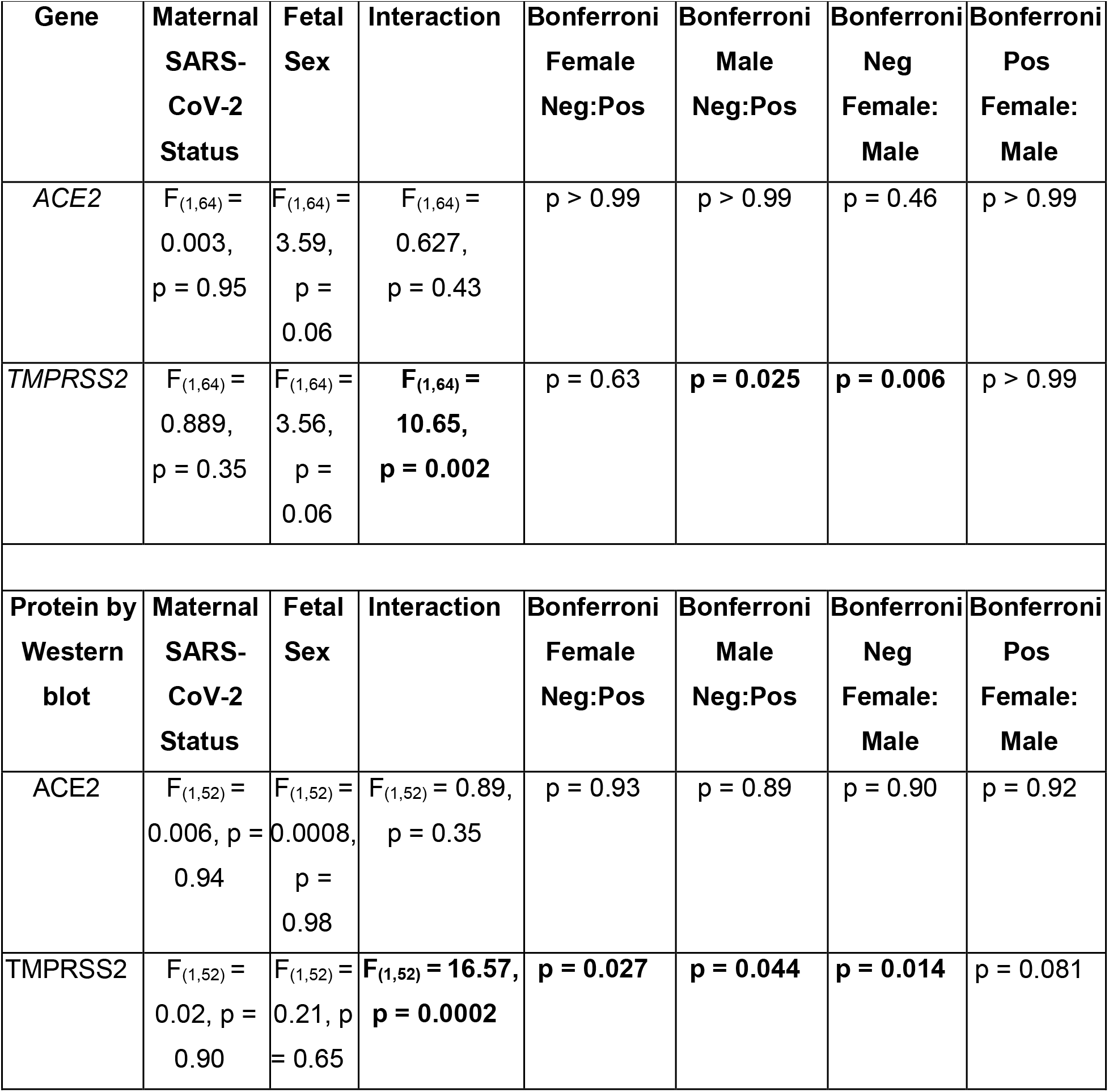
Two-way ANOVA analyses of ACE2 and TMPRSS2 gene and protein expression results. Two-way ANOVA followed by Bonferonni’s post-hoc analyses were performed to determine significance. All main and interaction effects for genes and proteins of interest are represented for both fetal males and fetal females in addition to all post-hoc analyses performed. Significant effects are indicated by bolded statistics. ”Neg” and “Pos” designate maternal SARS-CoV-2 status.

**Figure 1.**
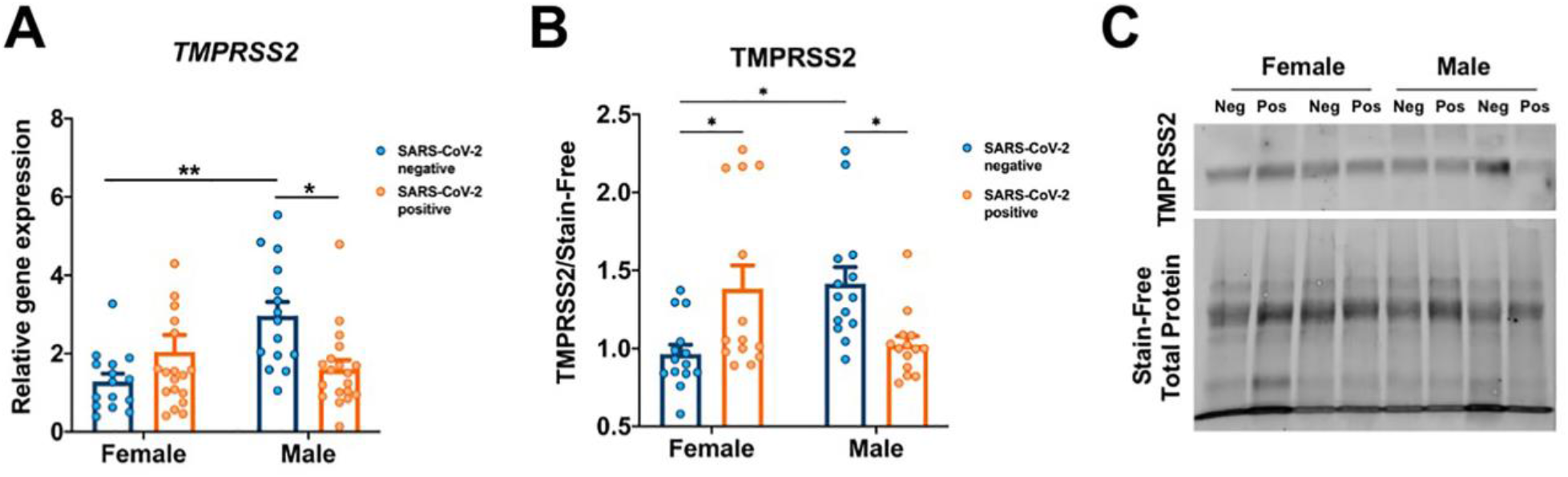
Sexually dimorphic placental TMPRSS2 expression in the setting of maternal SARS-CoV-2 infection. **A**. *TMPRSS2* gene expression demonstrates a significant interaction effect of fetal sex and maternal SARS-CoV-2 infection (interaction p-value: 0.002). *TMPRSS2* expression is elevated in male placentas of SARS-CoV-2 negative mothers compared to female placentas, suggesting higher baseline *TMPRSS2* expression in males. In the setting of maternal SARS-CoV-2 infection, *TMPRSS2* gene expression is significantly reduced in male placentas, with no significant effect of maternal SARS-CoV-2 infection in females. All expression levels are relative to reference gene *YWHAZ*. **B**. TMPRSS2 protein expression by Western blot analysis also demonstrates a significant interaction between fetal sex and maternal SARS-CoV-2 infection (interaction p-value: <0.001). In SARS-CoV-2 negative mothers, TMPRSS2 is elevated in male compared to female placentas. In SARS-CoV-2 positive mothers, TMPRSS2 is significantly reduced in male placentas and increased in female placentas relative to sex-matched controls. **P<0.01, *P<0.05. Data analyzed by 2-way ANOVA with Bonferroni’s post-hoc testing. **C**. Representative Western blots showing TMPRSS2 expression between males and females from pregnancies with and without exposure to maternal SARS-CoV-2 infection. “Neg” and “Pos” designate maternal SARS-CoV-2 status.

### No differences in ACE2 levels were observed by fetal sex or maternal SARS-CoV-2 infection

Relative to reference gene *YWHAZ*, there was no significant difference in placental *ACE2* gene expression in placental homogenates by fetal sex or in the presence of maternal SARS-CoV-2 infection (2-way ANOVA, fetal sex: P=0.06; SARS-CoV-2 exposure: P=0.95, interaction: P=0.43, Fig. 2A). These findings were confirmed by Western blot, which demonstrated no difference in ACE2 protein levels between groups (2-way ANOVA, fetal sex: P=0.98; SARS-CoV-2 exposure: P=0.94, interaction: P=0.35, Fig. 2B-C).

**Figure 2.**
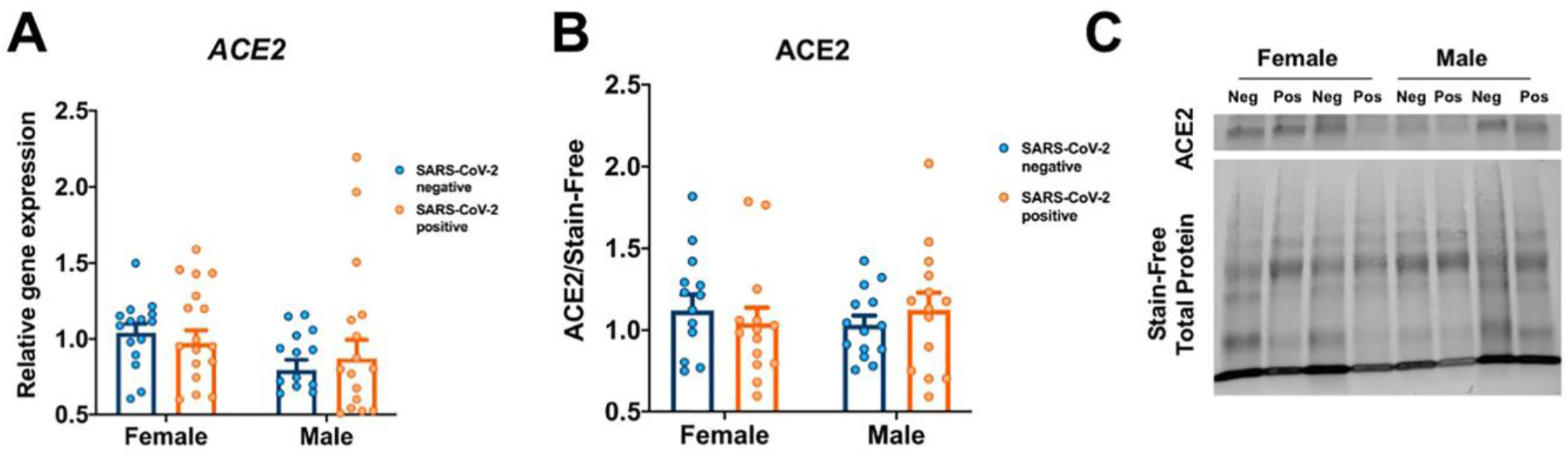
ACE2 expression does not vary by fetal sex or maternal SARS-CoV-2 status. **A**. *ACE2* gene expression by fetal sex and maternal SARS-CoV-2 infection, relative to female negative controls, demonstrating no significant differences between groups. All expression levels are relative to reference gene *YWHAZ*. **B**. No significant differences were observed in ACE2 protein expression by Western blot analysis. Data analyzed by 2-way ANOVA with Bonferroni’s post-hoc testing. **C**. Representative Western blots showing ACE2 expression between males and females from pregnancies with and without exposure to maternal SARS-CoV-2 infection.

### TMPRSS2 was significantly correlated with ACE2 expression in male placentas exposed to maternal SARS-CoV-2

We next sought to identify whether *ACE2* and *TMPRSS2* gene expression levels were significantly correlated within the same placenta in a sex-dependent fashion. Placental *TMPRSS2* expression was significantly correlated with *ACE2* expression in males (Spearman’s ρ=0.42, P=0.01) but not females (ρ=0.14, P=0.42), as demonstrated in Figure 3A. This relationship was driven by a significant correlation in male placentas exposed to SARS-CoV-2 (ρ=0.54, P=0.02, Fig. 3B) while SARS-CoV-2 unexposed male placentas had no significant correlation between ACE2 and TMPRSS2 gene expression (Fig. 3B). Placental *TMPRSS2* and *ACE2* expression levels were not significantly correlated in either female cases or controls (Fig. 3C).

**Figure 3.**
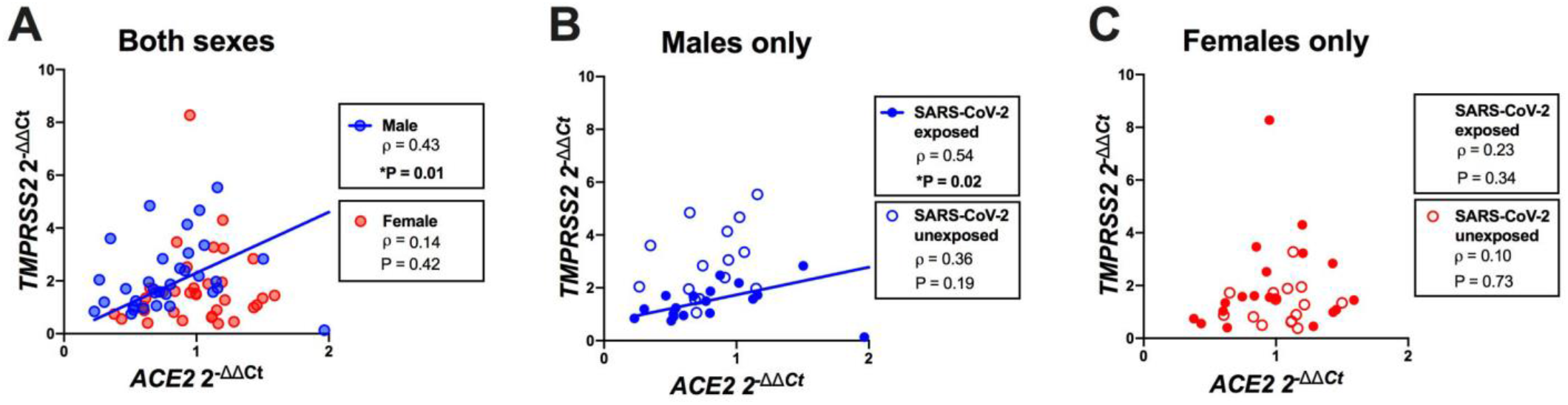
Correlation of *ACE2* and *TMPRSS2* expression within the same placenta. **A**. *TMPRSS2* is significantly correlated with *ACE2* in fetal male placentas (ρ=0.43, p=0.01, n=34) but not females (ρ=0.14, p=0.42, n=34). **B**. In fetal male placentas, *TMPRSS2* was significantly correlated with *ACE2* in SARS-CoV-2 exposed pregnancies (ρ=0.54, p=0.02, n=19) but not controls (ρ=0.36, p=0.19, n=15). **C**. There was no significant correlation between *ACE2* and *TMPRSS2* expression in fetal female placentas in either cases (ρ=0.23, p=0.34, n=19) or controls (ρ=0.10, p=0.73, n=15). All ddCt values for *TMPRSS2* and *ACE2* expression are normalized to female negative controls.

## Discussion

Here we describe for the first time baseline sex differences in TMPRSS2 gene and protein expression in the human placenta, and sexually dimorphic placental expression of TMPRSS2 gene and protein in the setting of maternal SARS-CoV-2 infection, even in the absence of placental infection. The consistent expression of TMPRSS2 transcript and protein in third trimester placentas using qRT-PCR and Western blotting suggest that standard immunohistochemical methods demonstrating weak or no expression of TMPRSS2 in the placenta [12,15] may lack the sensitivity to reliably detect TMPRSS2, which is more lowly expressed than ACE2. We demonstrate no changes in placental ACE2 gene or protein expression in the setting of fetal sex nor maternal SARS-CoV-2 infection. Maternal SARS-CoV-2 infection is associated with increased correlation between *ACE2* and *TMPRSS2* transcript levels in male placentas only. The impact of these sex-specific placental changes in canonical cell entry mediators for SARS-CoV-2 is not yet known, given the relative rarity of placental infection and in utero transmission of SARS-CoV-2 and lack of reported sex disaggregated data in these areas [2–4]. Reporting the sex differences described here represents the first essential step toward a more nuanced understanding of how fetal sex may influence vulnerability or resilience to placental and fetal infection with SARS-CoV-2. The finding of sex differences in TMPRSS2 expression may have implications for understanding sex differences in COVID-19 symptomatology [48–50], and for elucidating sex differences in diseases for which TMPRSS2 is implicated in the pathogenesis, such as H1N1 influenza and prostate adenocarcinoma [51,52].

TMPRSS2, which is highly expressed in prostate and lung tissue, among other sites, enables cellular infection with SARS-CoV-2 through priming of the Spike protein to facilitate binding the ACE2 receptor [53,54]. The strong regulation of TMPRSS2 by androgens has inspired the hypothesis that the male predominance in COVID-19 disease might be explained in part by increased TMPRSS2 levels [33,36]. A large scale cohort study investigating TMPRSS2 variants in an Italian population with high COVID-19 mortality rates, and male to female case ratio of 1.75, identified increased population frequency of a rare allele associated with increased TMPRSS2 expression [55]. Based on observational data that men on androgen-deprivation therapy for cancer risk reduction are at lower risk for acquiring SARS-CoV-2 [56], both androgen-deprivation therapy and agents blocking TMPRSS2 activity have been considered as therapeutic options for SARS-CoV-2 [24,54].

Consistent with evidence in support of higher TMPRSS2 expression in other male tissues, we identified higher TMPRSS2 levels in placentas of male compared to female offspring. Whether the observed higher baseline levels of TMPRSS2 in male placenta observed in our study represent increased vulnerability to placental infection is intriguing, but cannot be assessed in our cohort, as we identified no cases of placental infection. We did, however, identify a unique sexually dimorphic response to maternal SARS-CoV-2 infection in placental TMPRSS2 levels at both the gene and protein level: in females, maternal SARS-CoV-2 is associated with higher TMPRSS2 expression, whereas in males, maternal SARS-CoV-2 is associated with reduced TMPRSS2 expression. This observation is particularly important given the known increased vulnerability of the male fetus to prenatal insults [57–59]. Whether lower TMPRSS2 levels in the presence of maternal SARS-CoV-2 infection reflect a protective compensatory process in the male placenta warrants further study. Recent data from the CDC demonstrate slightly lower rates of vertical transmission in male neonates compared to female, with females accounting for 50.8% of SARS-CoV-2 diagnosed at birth, males accounting for 47.5% of cases, and 1.7% of SARS-CoV-2 diagnosed perinatally occurring in neonates with unknown/not recorded sex at birth (overall N=59) [60]. Given the small number of reported cases of perinatal transmission, future large cohorts reporting on vertical and perinatal transmission should do so in a sex-disaggregated fashion to further our understanding of the potential for male vulnerability to or protection from SARS-CoV-2 vertical transmission.

We identified no significant effect of fetal sex or maternal SARS-CoV-2 infection on placental ACE2 gene or protein expression. As part of the renin-angiotensin-aldosterone system (RAAS) that regulates fluid homeostasis, the ACE2 enzyme is present in various cell types throughout the body, including the lung and respiratory tract [62]. Sex steroids - androgens and estrogens - influence the RAAS system [63,64] and sex-specific differences in ACE2 expression render males more vulnerable to renal and cardiovascular disease, with higher levels considered more detrimental [65–67]. Evidence suggests that sex steroids can influence ACE2 expression in the respiratory tract, and that higher ACE2 levels may be causally related to increased COVID-19 severity [67–71]. However, studies investigating ACE2 expression in respiratory tract tissues by sex have not consistently shown higher expression in males, highlighting that levels of ACE2 may not fully explain the observed male bias in severe COVID-19 disease [55,71–74]. While our data and several other studies demonstrate robust expression of ACE2 in the human placenta that may vary by gestational age [12,15,17–19,75,76], few studies have evaluated sex-specific placental ACE2 expression, and these have reported conflicting results [78,79]. We are not aware of any studies to date that have evaluated sex differences in placental ACE2 expression in the setting of maternal SARS-CoV-2 infection. A study of maternal decidual explant cultures reported that secreted *ACE2* mRNA was significantly increased in 48-hour explant cultures from mothers carrying a female fetus [78], while evaluations of rat placenta have found reduced *ACE2* expression in the setting of maternal protein restriction and dexamethasone administration [79,80], but no sex differences in this regard. Our data suggest that ACE2 expression in placental tissue does not differ significantly by offspring sex and is not significantly affected by maternal SARS-CoV-2 infection. Based on this finding, differences in placental ACE2 levels likely do not represent a source of sex-specific vulnerability to placental infection or vertical transmission.

We found that *ACE2* and *TMPRSS2* transcript levels are highly correlated in male placentas in the setting of maternal SARS-CoV-2 infection, but not in uninfected male pregnancies, nor in female placentas regardless of maternal infection status. Expression profiling of ACE2 and TMPRSS2 in human tissues has shown a strong positive correlation across multiple cell lines and tissue types [82], with authors postulating that any intervention or mechanism targeting one gene may affect expression of the other. The same study found that ACE2 and TMPRSS2 tend to be co-regulated by factors such as obesity, diabetes, viral infections, and androgens [82]. TMPRSS2 is known to be regulated by androgen/androgen receptor signaling in prostate cancer [83], and a recent study identified positive correlations between androgen receptor and ACE2 expression in multiple tissues [38]. Further investigation into the interplay between maternal SARS-CoV-2 infection and androgen signaling in male placenta may yield important insight into the placental response to maternal infection with SARS-CoV-2 or other viral pathogens.

Our histopathological data from term placentas demonstrating low expression of TMPRSS2 relative to ACE2 and physically distant expression of these two cell entry mediators [12,15] suggest that the increased correlation of ACE2 and TMPRSS2 expression observed here in male SARS-CoV-2-exposed placentas may not represent increased co-localization of ACE2 and TMPRSS2. Both negligible co-transcription of placental ACE2 and TMPRSS2 across gestation by nearly all placental cells [17] and the relatively low expression of TMPRSS 2 relative to ACE2 in full term placenta [12] have been cited as potential mechanisms protecting against placental SARS-CoV-2 infection. It remains unclear, however, whether enhanced synchronicity in ACE2/TMPRSS2 expression patterns represents an increased placental vulnerability to infection.

Strengths of our study include a relatively large sample size of pregnancies with SARS-CoV-2 infection, including 58% with symptomatic illness and 8% with severe or critical disease. Another strength of this cohort is the identification and inclusion of a contemporaneously-enrolled control population. To control for heterogeneity and regional differences in the placenta, we used four biological replicates with equal representation of fetal and maternal surfaces. Our study is limited by the timing of SARS-CoV-2 infection in pregnancy, which largely occurred in the third trimester. The effects of first or second trimester infection on *ACE2* and *TMPRSS2* expression should be evaluated, as infection earlier in pregnancy could be associated with increased risk of vertical transmission. The impact of sex differences in placental patterns of ACE2 and TMPRSS2 expression will need to be evaluated in large cohorts with the ability to link biological data with disease outcomes of interest such as vertical transmission.

These data demonstrate striking sex differences in placental patterns of SARS-CoV-2 cell entry mediators in the setting of maternal SARS-CoV-2 infection. Moreover, the baseline sex differences in placental TMPRSS2, and the sexually dimorphic response of TMPRSS2 to maternal infection even in the absence of placental infection, suggest a potential role for TMPRSS2 in the placenta beyond SARS-CoV-2 entry that may be affected by fetal sex or maternal exposures. The findings presented here demonstrate the importance of reporting sex disaggregated placental and neonatal outcomes data in the setting of maternal SARS-CoV-2 infection. These data also elucidate potential placental defenses against viral infection with SARS-CoV-2, including altered TMPRSS2 expression in the presence of maternal SARS-CoV-2 infection.

## Abbreviations

(ACE2): Angiotensin-Converting Enzyme 2
(TMPRSS2): Type II Transmembrane Serine Protease
(YWHAZ): Tyrosine 3-Monooxygenase/Tryptophan 5-Monooxygenase Activation Protein Zeta
(RAAS): renin-angiotensin-aldosterone system

